# Development and deployment of improved *Anopheles gambiae* s.l. field surveillance by adaptive spatial sampling design

**DOI:** 10.1101/2023.06.16.545360

**Authors:** Gabriel M. Monteiro, Luc S. Djogbénou, Martin J. Donnelly, Luigi Sedda

## Abstract

Accurate assessments of vector occurrence and abundance, particularly in widespread vector-borne diseases such as malaria, is essential for efficient deployment of disease surveillance and control interventions. This study emphasizes the need for flexible spatial sampling designs that can capture the dynamic relationships between disease vector species and the environment. Although previous studies have examined the benefits of adaptive sampling for disease hotspot identification (mostly by simulations), limited research has been conducted on field surveillance of malaria vectors. Here, an adaptive spatial sampling design targeting potential and uncertain *An. gambiae* hotspots, a major malaria vector in sub-Saharan Africa, is presented. The first phase of the proposed design involved ecological zone delineation and a proportional lattice with close pairs sampling design to maximise spatial coverage, representativeness of ecological zones and vector spatial autocorrelation (by the employment of close pairs). In the second phase, a spatial adaptive sampling design targeted high-risk areas with the largest uncertainty. For the second phase, the sample size was reduced compared to the first phase, but predictions improved for out-of-sample and training data. However, the overall model uncertainty increased, highlighting the trade-off in multi-criteria adaptive sampling designs. It is important that future research focuses on these trade-offs to reduce the timescale for effective malaria control and elimination efforts.

## Introduction

The Global Vector Control Response (GVCR) was unanimously adopted by the World Health Assembly in May 2017 to tackle the stall in malaria elimination progresses (World Health Organization and UNICEF 2017). Since then, WHO has engaged in a programme to roll out the GVCR in all regions. GVCR is based on four pillars for Vector-Borne Disease control: intersectoral collaboration; community engagement; monitoring, surveillance and evaluation; and integration of tools and approaches, supported by novel and innovative research.

The four pillars for malaria control and elimination are reliant upon disease risk in estimated in time and space, which must be routinely reviewed to inform, tailored malaria control strategies (Thawer, Golumbeanu et al. 2022) and the impacts of these interventions (Shrestha, McCulloch et al. 2022). A key component in the definition of the disease risk is the deployment of optimal entomological surveillance frameworks, where optimal can be defined as the strategy providing the maximum accuracy in the investigated mosquito process given limited resources(Sedda, Lucas et al. 2019). An Entomological Adaptive Sampling Framework (EASF) adjusts the entomological sampling strategy across time and space and enables the capture of spatially heterogeneous changes in entomological and disease transmission dynamics, and guide programmatic and strategic decisions (Obsomer, Titeux et al. 2013). Within this framework, cost-effectiveness cannot only be dependent on accessibility of the survey locations but also on the robustness and quality and quantity of information of its obtained data (Koenraadt 2021), which is the priority for any EASF.

For an entomological survey to provide unbiased information on species occurrence and abundance while detecting the spatio-temporal relationships between species and environment, a degree of spatial spread of sample locations including areas where these relationships are weaker is required. Malaria, and other vector-borne disease systems, are complex due to spatial and temporal heterogeneity (Kabaghe, Chipeta et al. 2017), and therefore require flexible spatial sampling designs to represent the full dynamicity of the disease system components. In absence of prior knowledge of the process under investigation (expert opinion, historical data), a spatially balanced design which spreads the survey locations as evenly as possible in the study area according to a specific sampling method or criteria, provides more uniform coverage over the study area than random sampling and decreases uncertainty in spatial interpolation (Gelfand, Sahu et al. 2012). Spatial balanced designs can be obtained by employing quasi-random methods (Sobol and Halton sequence) or distance-based designs commonly known as space filling designs (Liu and Vanhatalo 2020). The major drawback of these designs, in the presence of prior information, is that locations are selected with identical or roughly equal probabilities. However, when previous data are available sampling can be based on a target function (often model-based). Unequal probability survey designs or adaptive designs allow selection of locations based on a specific criteria, and sample locations are selected based on different probabilities of appearing in the sample (Brown, Salehi et al. 2013).

With spatially correlated data, adaptive sampling and sequential sampling, often associated with stratified sampling can increase the information content and provide a more efficient estimation of the vector or its disease (Lazaro, Sese et al. 2021). Often these methods target locations with high spatial uncertainty or disease hotspots (Case, Young et al. 2022).

In the field study reported here, a spatially balanced design coupled with adaptive sampling was employed for entomological surveillance in the south west region of Benin (West Africa). We evaluate the efficacy of spatial adaptive sampling designs for vector surveillance. Strategy and model performances and uncertainty changes for the primary malaria vector *Anopheles gambiae* s.l. (simply *Anopheles gambiae* thereafter) are presented.

## Methods

### Study area

The study site encompasses the provinces of Athiémé, Bopa, Comè, Grand Popo, Houeyogbé, Kpomassè, Ouidah and Sè in the region of Atlantique (South West Benin). This region expands from the south west coast to 30km inland and from 0 m to 70m above sea level. The average temperature is 28.9°C, average relative humidity is 76% with average annual rainfall of 190mm. The wet season is characterized by abundant rains between April and July, with a lower amount of rain from September to October (Boton, Fangninou et al. 2019). The transmission of malaria is perennial in the region where malaria infection is mesoendemic or hyperendemic (malaria prevalence between 40 and 60%) (Damien, Djènontin et al. 2010)

### Spatial sampling design

Entomological surveillance was carried out in two phases. The first phase (Phase I) was carried in 2018 and based in a spatially balanced sampling design (lattice with close pairs) as described in (Sedda, Lucas et al. 2019). The following entomological surveillance in 2021 (Phase II) was based on a spatial adaptive sampling design (Chipeta, Terlouw et al. 2016), targeting *Anopheles gambiae* highest intensities, with highest uncertainties, based on the prediction results from Phase I.

An adaptive spatial sampling scheme entails selecting a number of sampling locations over a sequence of sampling times through probabilistic models. At each step, the data analysis is executed using an enriched sample data set. The inherent spatial autocorrelation is explicitly considered to improve prediction and inference. The implementation of the adaptive sampling design required four preliminary decisions: a) an initial model to fit the data from Phase I; b) a batch size as the number of new locations to add to the first phase sampling design; c) a utility function to rank unobserved locations to target; and d) removal of low information locations from Phase I to satisfy limited availability of resources.

The Phase I *An. gambiae* counts per week (*w*) and household (*h*), *Y_w,h_*, were fitted by employing a Poisson generalised linear mixed model with independent and identically distributed spatiotemporal random fields:

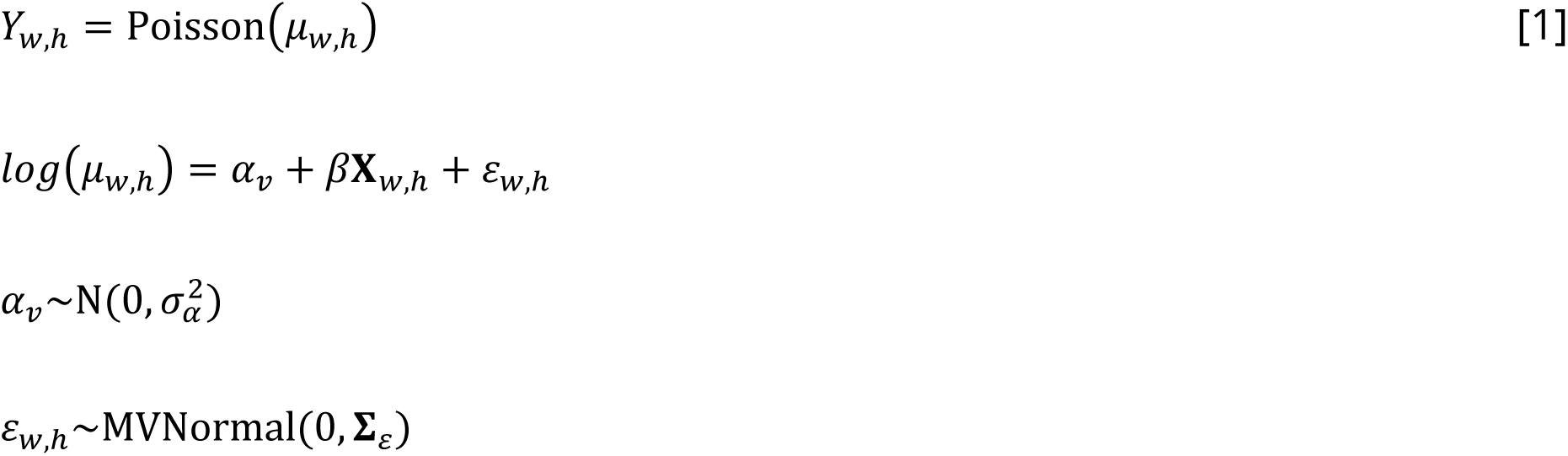

where *α* is a group-level random effect normally distributed with mean equals to 0 and variance *σ*^2^, with group identified by the location id, *v*; *β* is a vector of coefficients for each explanatory variable contained in the matrix X, *ε* is the spatiotemporal random Gaussian field with covariance matrix Σ_ε_ constrained by a Matern covariance function. This model is implemented by using sdmTMB package in R-cran software (Anderson, Ward et al. 2022). The spatiotemporal random filed is approximated through a triangulated mesh with minimum gap of 1 km.

In this modelling context, a risk map is equivalent to mapping detection probabilities for *An. gambiae* (Aarts, Fieberg et al. 2012).

The adaptive batch was 15 new locations (half of the current ones) as constrained by available resources. The utility function is based on the predicted intensity of *An. gambiae* and therefore 15 locations were selected by ranking the largest posterior predicted risk and within them the ones with largest uncertainty. This strategy can lead to high sampling bias and can fail in prioritising locations with new information because not exploring areas with low intensity in the search space (Case, Young et al. 2022). However, the focus of this work was to delineate *An. gambiae* hotspots in known and unknown locations since Phase I was run on a relatively short period.

Finally, a jackknife approach was implemented to discard locations from Phase I containing limited information, i.e. it has limited impact on the uncertainty in predictions when dropped (Wang, Moe et al. 2020).

### Environmental variables

*Anopheles gambiae* counts were inferred and predicted based on co-collected *An. funestus* s.l. (*An. funestus* thereafter) and eco-strata created in (Sedda, Lucas et al. 2019). The use of *An. funestus* as predictor can provide information around co-occurrence of the two mosquito species and therefore a more precise quantification of malaria risk (Djènontin, Bio-Bangana et al. 2010). No other ecological variables were used because the eco-strata is a construct of several environmental satellite data obtained from open sources: land cover classification at 30m resolution obtained from GlobeLand30 for the year 2010 and containing 10 classes (Jun, Ban et al. 2014); elevation from the NASA Shuttle Radar Topographic Mission (SRTM) at 90m and released in 2008 (Jarvis, Reuter et al. 2008); bioclim precipitation as average annual precipitation from 1970 to 2000 at 30 arcseconds (1 km ca.)(Fick and Hijmans 2017); soil information at 30 arcseconds from the FAO harmonized world soil database V1.2 (Fischer, Nachtergaele et al. 2008); and finally, monthly MODerate-resolution Imaging Spectroradiometer (MODIS) satellite products for temperature, enhanced vegetation index (EVI) and the ratio of Actual to Potential Evapotranspiration (ET) were obtained from Oxford University Research Archive at 0.05 degrees (5 km ca.) (Seddon, Macias-Fauria et al. 2016). For each of the MODIS variables, the mean, amplitude and variance for the entire 2000-2013 period were calculated. We identified four eco-strata: 1, cultivated land/grassland; 2, forest/shrubland; 3, wetland and water bodies; and 4 urban.

Given these environmental variables the glmm model above can be detailed as:

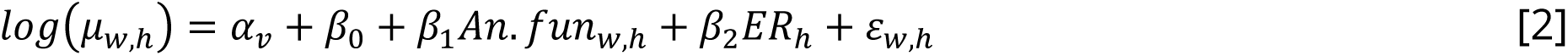

where An. fun are the counts of *An. funestus*, ER is the eco-strata id, and the *β* are the intercept (_0_), and coefficients for *An. funestus* (_1_) and each eco-strata (_2_).

### Entomological collection

The design of the first entomological survey of 30 locations, based on a spatial balanced design, has been described by (Sedda, Lucas et al. 2019).

After employing the adaptive sampling design, entomological collections were carried out from October to November 2021 in seventeen locations (see results) for a total of 68 houses (four houses per location). Houses were randomly selected for mosquito collection. Mosquitoes were collected during 4 consecutive nights every week for 4 weeks, using CDC light traps (Model 512, John W. Hock Company, Gainesville, FL). For each location, a member of the community was trained in the use of the traps and sampling protocol. The order of house collection was randomised before each collection to remove systematic biases. Traps were positioned at a height of 1.5 m in sleeping rooms and operated on 12V battery power from 8 p.m. to 6 a.m. Each morning, mosquitoes were collected, stored in ziplock bags containing silica gel and transported immediately to the laboratory of the Tropical Infectious Diseases Research Center of the University of Abomey-Calavi located at the Institut Regional de Santé Publique of Ouidah for identification and analyses.

### Model and prediction performance

Model parameters uncertainty were provided by 95% credible intervals (a more robust measure than confidence intervals when comparisons are made between models with different sample size), and compared between Phase I and Phase II and Phase I and II combined. Comparison was by estimation of credible intervals probability of overlapping (Kruschke 2013).

Predictions at both Phase I and Phase II and for the overall Phase I and II were obtained on a prediction grid with 5 km spatial resolution covering the all study area (longitude from 1.5 to 2.1 E degrees and latitude from 6 to 7 N degrees). Uncertainty is obtained by calculating the point grid standard deviation of the *An. gambiae* estimations from 999 simulations of the respective fitting models, with parameters draws from their posterior distribution via Maximum Likelihood Estimation (Anderson, Ward et al. 2022).

The Leave-One-Out Cross-Validation (Fuhg, Fau et al. 2020) was employed to evaluate out of sample prediction errors by removing one record from the training set and predicting the value of *An. gambiae* in the removed record (a record is one of the house-location-week *An. gambiae* catch). The procedure was repeated 100 times and an average squared error measured.

Finally, model explained variance and training data prediction error were calculated from r-squared (Gelman, Goodrich et al. 2019) and the root mean squared error (RMSE) (Rocha, Groen et al. 2021) statistics respectively. A lower RMSE value indicates that the predicted values are closer to the actual values, while a higher RMSE value indicates that the predicted values are further away from the actual values.

### Ethical approval

Secondary data analyses have been approved by the Faculty of Health and Medicine Research Ethics Committee at Lancaster University (UK) with reference number FHMREC20173.

## Results

The *An. gambiae* counts obtained during Phase I were used to produce an *Anopheles gambiae* risk map (Figure 1a).

**Figure 1.**
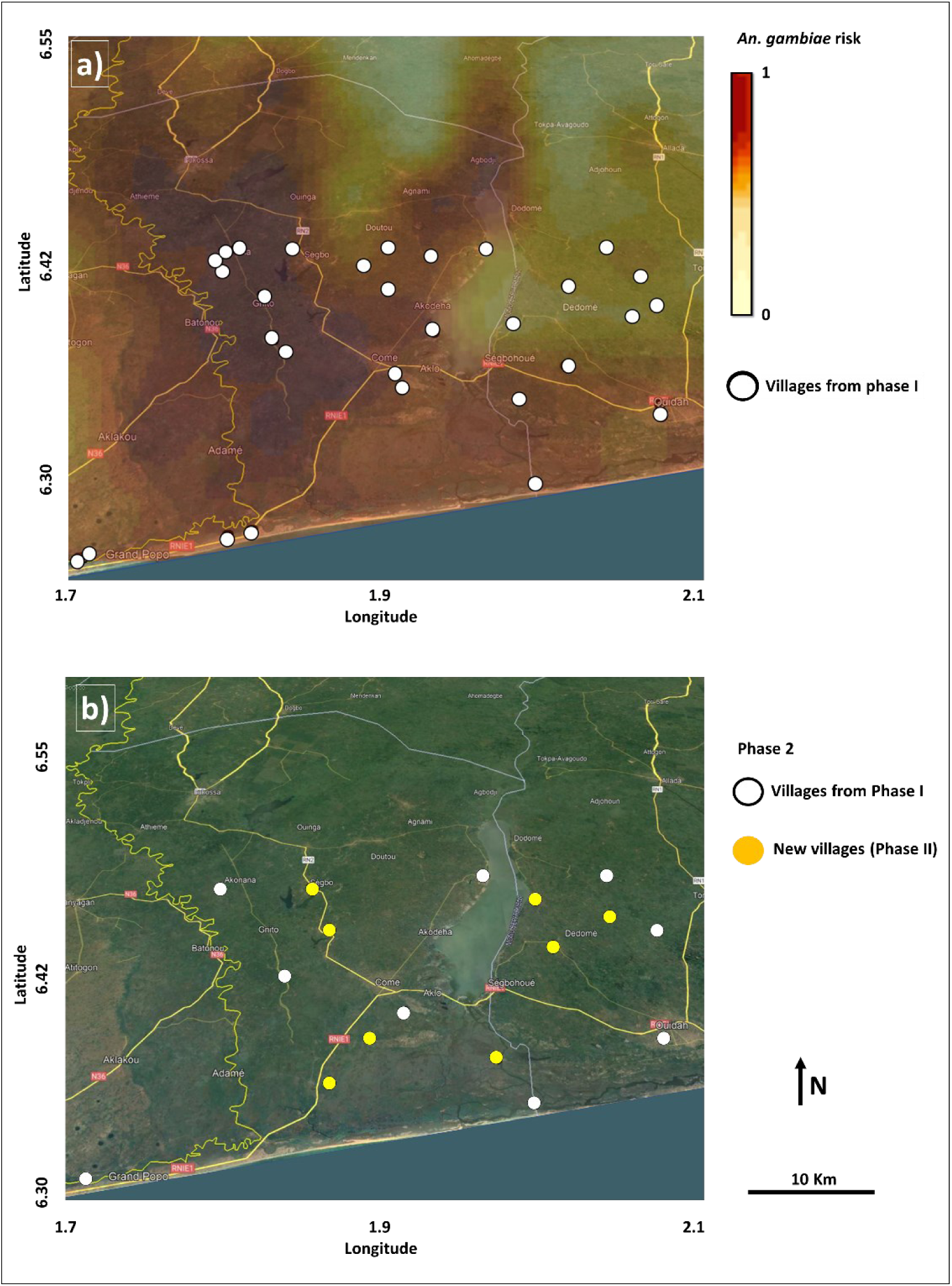
(a) *Anopheles gambiae* risk map (darker colors higher risk) with dots in background showing sampling locations in Phase I. (b) Based on the *An. gambiae* risk, Phase II surveillance used a mixture of previous locations (white dots) and new locations (yellow dots). Background: Google Earth Image@2021 (Image Landsat/Copernicus; CNES/Airbus; Maxar Technologies. Data: SIO, NOAA, U.S. Navy, NCA, GEBCO).

The spatial adaptive sampling allocated 15 locations with high *An. gambiae* risk and high uncertainty, 8 in new locations and 7 overlapping with Phase I locations. The jackkinfe loss of information criteria method removed 21 locations from Phase I. This led to a Phase II on 17 locations (8 new and 9 from Phase I) (Figure 1b).

Overall a larger amount of *An. gambiae* were caught in Phase I compared to Phase II. On the contrary, an average larger amount of *An. funestus* was caught in Phase II than in Phase I (Table 1). Despite this, the two species show a significant positive association between them (Table 2).

**Table 1.** Summary statistics for catches of *An. gambiae* and *An. funestus* by sampling phase. Sampling effort is defined by the product of the number of houses in each location, the number of locations and the number of weeks of trapping (720 from 4 houses, 30 locations and 6 weeks; 272 from 4 houses, 17 locations and 4 weeks).

| Species/Phase | Mean | Median | Min | Max | SD | Sampling Effort |
| --- | --- | --- | --- | --- | --- | --- |
| *An. gambiae* I | 4.80 | 1 | 0 | 213 | 12.45 | 720 |
| *An. gambiae* II | 1.77 | 0 | 0 | 34 | 4.86 | 272 |
| *An. gambiae* I+II | 3.97 | 0 | 0 | 213 | 10.98 | 992 |
| *An. funestus* I | 0.21 | 0 | 0 | 23 | 1.28 | 720 |
| *An. funestus* II | 0.56 | 0 | 0 | 13 | 1.55 | 272 |
| *An. funestus* I+II | 0.31 | 0 | 0 | 23 | 1.37 | 992 |

**Table 2.**
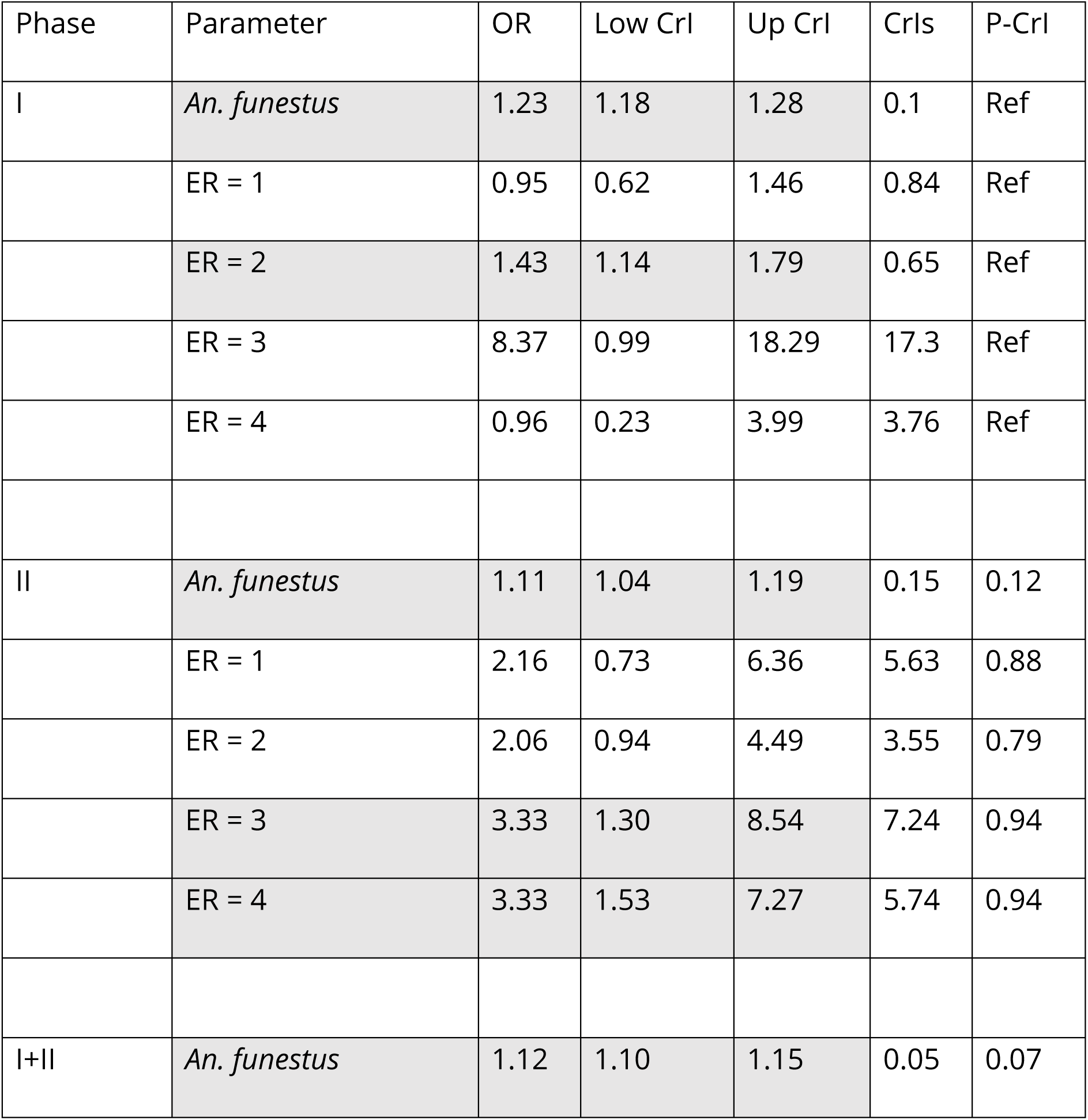
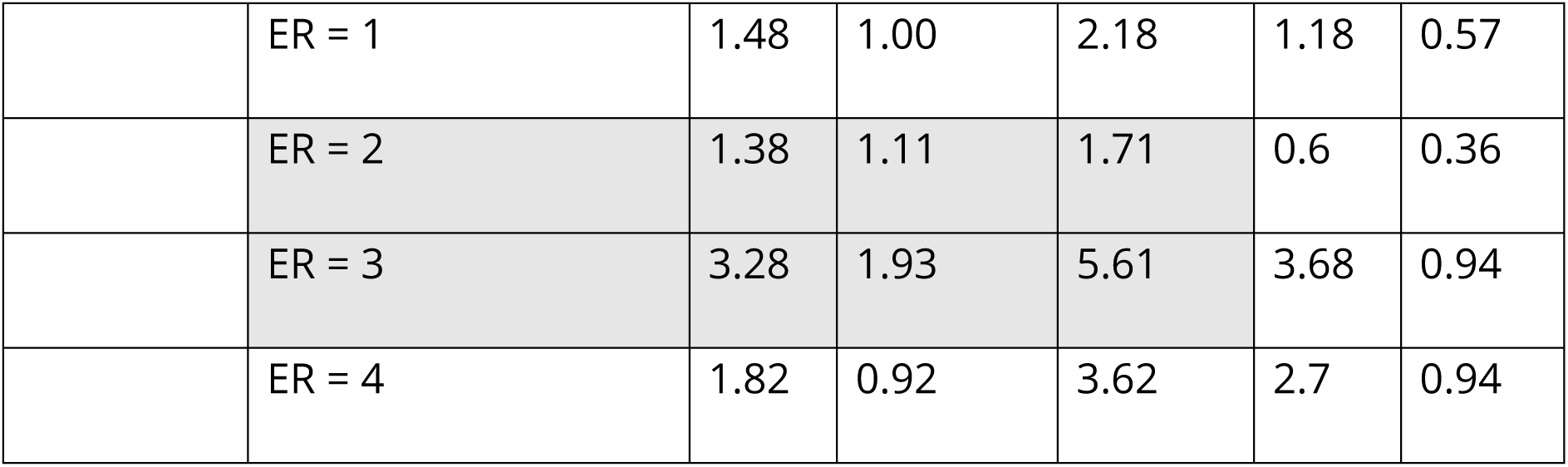
Poisson generalised linear model coefficients for fixed effects by modelling *An. gambiae* counts per week and household. OR, odd ratios; Low CrI, lower 95% credible interval; Up CrI, upper 95% credible interval; CrIs, credible interval size; P-CrI probability of overlap between the credible intervals of Phase II vs Phase I and Phase I+2 vs Phase I. ER ids: 1, cultivated land/grassland type; 2, forest/shrubland-type; 3, wetland and water bodies; and 4 urban.

| Phase | Parameter | OR | Low CrI | Up CrI | CrIs | P-CrI |
| --- | --- | --- | --- | --- | --- | --- |
| I | *An. funestus* | 1.23 | 1.18 | 1.28 | 0.1 | Ref |
|  | ER = 1 | 0.95 | 0.62 | 1.46 | 0.84 | Ref |
|  | ER = 2 | 1.43 | 1.14 | 1.79 | 0.65 | Ref |
|  | ER = 3 | 8.37 | 0.99 | 18.29 | 17.3 | Ref |
|  | ER = 4 | 0.96 | 0.23 | 3.99 | 3.76 | Ref |
| II | *An. funestus* | 1.11 | 1.04 | 1.19 | 0.15 | 0.12 |
|  | ER = 1 | 2.16 | 0.73 | 6.36 | 5.63 | 0.88 |
|  | ER = 2 | 2.06 | 0.94 | 4.49 | 3.55 | 0.79 |
|  | ER = 3 | 3.33 | 1.30 | 8.54 | 7.24 | 0.94 |
|  | ER = 4 | 3.33 | 1.53 | 7.27 | 5.74 | 0.94 |
| I+II | *An. funestus* | 1.12 | 1.10 | 1.15 | 0.05 | 0.07 |
|  | ER = 1 | 1.48 | 1.00 | 2.18 | 1.18 | 0.57 |
|  | ER = 2 | 1.38 | 1.11 | 1.71 | 0.6 | 0.36 |
|  | ER = 3 | 3.28 | 1.93 | 5.61 | 3.68 | 0.94 |
|  | ER = 4 | 1.82 | 0.92 | 3.62 | 2.7 | 0.94 |

**Table 3.** Model performance. RMSE, root mean squared error.

| Model | Variance spatio-temporal random effect (95% confidence intervals) | Leave-one-out cross validation error | Overall uncertainty | RMSE | R-squared |
| --- | --- | --- | --- | --- | --- |
| Phase I | 5.84 (3.19-10.07) | 0.16 | 0.19 | 1.27 | 0.16 |
| Phase I+II | 4.51 (3.30-6.16) | 0.12 | 0.23 | 1.11 | 0.27 |

When comparing Phase I and Phase I+II, all significant predictors for *An. gambiae* (*An. funestus* and forest/shrubland-type) had narrower credible intervals in combined data for Phase I and Phase II than in Phase I (probability of overlapping lower than 0.5) (Table 2). In both Phase I and Phase I+II, increased *An. funestus* catches and presence of mixed forest/shrubland eco-strata increased the likelihood to catch *An. gambiae* with mixed forest/shrubland eco-strata exerting a stronger effect than *An. funestus* (Table 2, OR values).

The model with Phase I+II data produced a lower variance of the spatio-temporal effect (meaning a larger amount of variance explained by the predictors than in the model of phase one) and in terms of *An. gambiae* estimation a lower error during cross-validation, a lower root mean squared error and a higher explained variance (R-squared). However, the general uncertainty in the area increased of around 21%.

Finally, Figure 2 shows the *An. gambiae* relative risk in Phase I and Phase I+II, the latter with general increased risk in most of the region but not in the north east.

**Figure 2.**
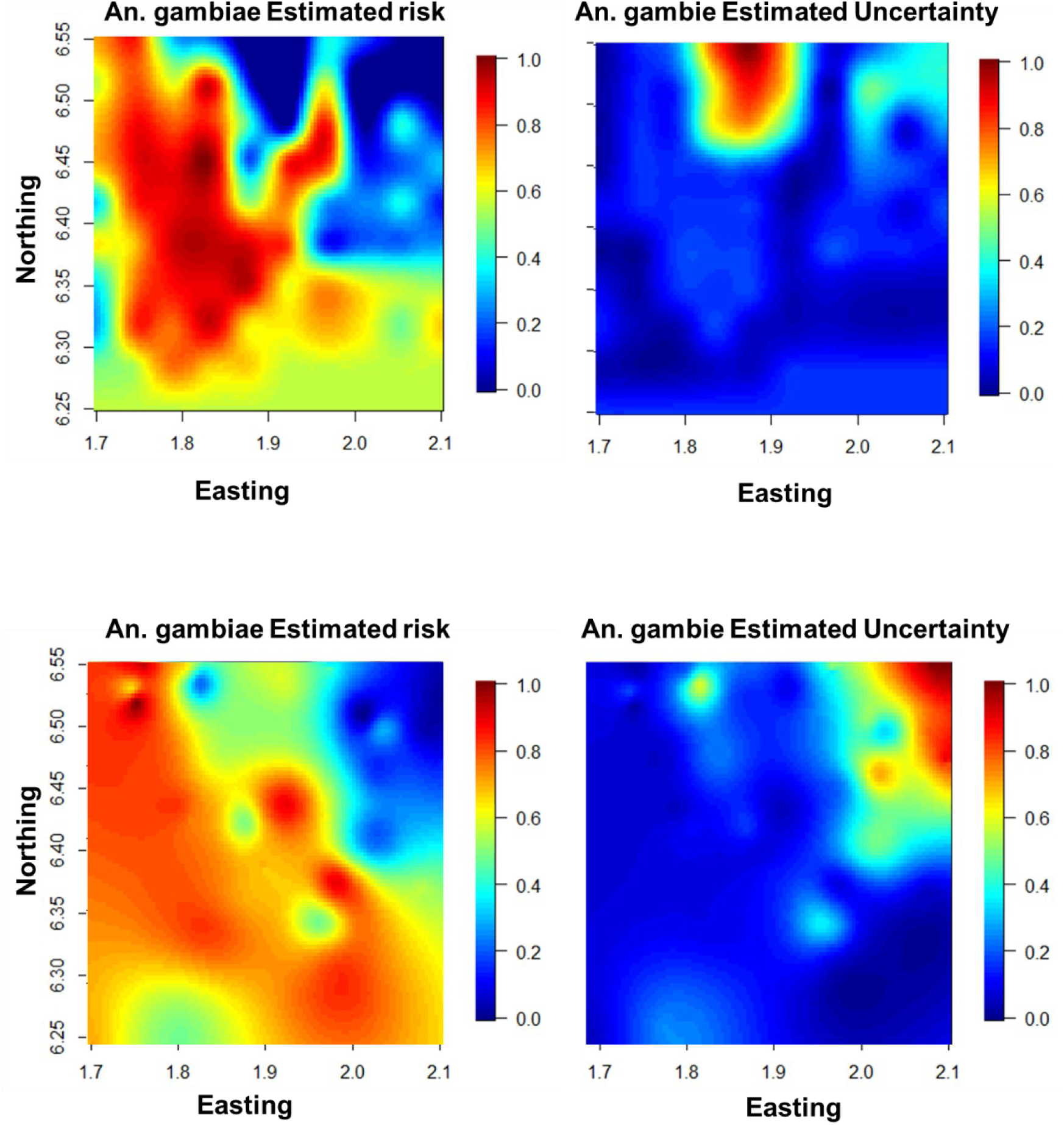
*Anopheles gambiae* relative risk and relative uncertainty in Phase I (top) and Phase I+II (bottom).

## Discussions

To provide scientifically grounded guidance to decision makers for effective, targeted malaria control and elimination, an accurate assessment of the spatio-temporal malaria transmission intensity that can take into account the spatio-temporal heterogeneities in both the vectors and parasites is essential. This will allow the design of tailored intervention strategies at different locations and time periods. Model-based sampling strategies, such as spatial adaptive sampling designs, are the best equipped tools to undercover these heterogeneities.

The present study was aimed at improving the definition of the most uncertain *An. gambiae* hotspots by targeting its largest and most uncertain risk. The research was carried out in two phases: a first phase where zone delineation (eco-strata) and a proportional lattice with close pairs sampling design was implemented; and a second phase that combined a spatial adaptive sampling design targeting high risk with highest uncertainty with information loss criteria method to add new locations and remove obsolete ones. The Phase II sample size was 56% of the sample size of Phase I. While out of sample and training data predictions improved with the additional added adaptive locations, the general uncertainty increased by a fifth of the initial one. In other words, the presented framework has an accurate expectation of the *An. gambiae* spatio-temporal heterogeneity (i.e. an accurate description of the average risk in the sampled area) although with larger margins of uncertainty when compared to the initial sampling. The used adaptive target criteria has certainly contributed to these results, since the utility function was not based on the global uncertainty. It is well known that uncertainty map can indicate where additional data would reduce the overall prediction error, and although other authors (Shrestha, McCulloch et al. 2022) suggest similar strategies as the one adopted in this work, we have shown that short-term targets (where short term is defined as low number of repeated sampling) are likely to improve only (partly or fully) the target characteristics of the model while potentially making worse other ones. Additional work on the effective capacity of multicriteria adaptive sampling design is then required with relation to the length and frequency of the surveillance.

Previous work concentrated on the benefits of spatial adaptive sampling strategies on the disease not the vector and often by simulation of artificial data (Case, Young et al. 2022). For example, Andrade-Pacheco and colleagues (Andrade-Pacheco, Rerolle et al. 2020) shown that a spatially adaptive sampling approach produced consistently superior accuracy for a generic disease hotspot classification over a random sampling approach, and could dramatically lower the resource requirements to conduct surveys whose goal is to detect disease hotspots. The only study found on field surveillance of malaria is the work of Kabaghe and others (Kabaghe, Chipeta et al. 2017) who applied adaptive sampling for malaria prevalence in an area in Malawi and identified areas where increased sampling effort increased overall predictive accuracy of the hotspots area.

However, their survey design targeted predictive high intensity areas of malaria prevalence only.

While the model presented herein shows a good robustness (RMSE around 1 which fits the common assumption of residual errors with 1 standard deviation), it is limited by the large time passing from Phase I and Phase II (almost 3 years) which may have contributed to the lower number of *An. gambiae* catches, and ignored potential interventions. In addition, this work does not compare with a control, e.g. a surveillance in the same region involving random sample, though the scientific community agree on the superiority of the adaptive or other model-based designs compared to random sampling. Finally, 21 locations from Phase I were removed due to minimal contribution to the overall information and satisfy the condition of limited available resources. However, this should be done carefully and future studies should evaluate the effect of removal sequentially instead of in batch to account for intra-location variation and seasonality effects at locations candidate for removal.

Future work will need to address the limitations of the presented study and answer other important questions for optimal interventions, such as the operational unit for treatment which will require the integration of the epidemiological component to the entomological one (Rebollo, Zoure et al. 2018); and on the epidemiological and entomological indicators that can consider the heterogeneity of parasites, vectors and environments (Liu, Liu et al. 2023). For any malaria elimination strategy, it will be necessary to balance the targeting of interventions to disease hotspot, with the need to correctly identify the vector and infestation status in the larger neighbourhood area to ascertain the risk of potential re-introduction (Huestis, Dao et al. 2019, Stolk, Blok et al. 2021, Case, Young et al. 2022) but also logistical constraints (e.g. human resources and cost). (Wang, Moe et al. 2020)

In conclusion, for malaria and many other infectious diseases, two conditions need to be met for the disease to be categorised as susceptible to eradication/elimination: first, there must be accurate diagnostic(s); and second, there must be effective intervention tool(s) able to remove the infection from the area. The latter will only succeed if appropriate sampling strategies are available. This work results provide new insights for the development of these strategies.

## Acknowledgments

The authors wish to thank the householders, villagers and local staff (Esdras Odjo Constantin Adoha and Pierre Sovegnon) for their contribution to the research.

## Contributors

MD and LS conceived the study. GM led field work and data curation. LS designed the algorithm and conducted the analyses. LS and LD acquired funding, administrated the project, and led the investigation. MD and LD provided supervision. GM and LS drafted the manuscript. All authors edited the manuscript and approved the final draft.

## REFERENCES

Aarts, G., J. Fieberg and J. Matthiopoulos (2012). “Comparative interpretation of count, presence-absence and point methods for species distribution models” Methods in Ecology and Evolution 3(1): 177–187.

Anderson, S. C., E. J. Ward, P. A. English and L. A. K. Barnett (2022). “sdmTMB: an R package for fast, flexible, and user-friendly generalized linear mixed effects models with spatial and spatiotemporal random fields.” bioRxiv: 2022.2003.2024.485545.

Andrade-Pacheco, R., F. Rerolle, J. Lemoine, L. Hernandez, A. Meite, L. Juziwelo, A. F. Bibaut, M. J. van der Laan, B. F. Arnold and H. J. W. Sturrock (2020). “Finding hotspots: development of an adaptive spatial sampling approach” Scientific Reports 10(1).

Boton, D. M., F. F. Fangninou, B. Xu and P. Houedegnon (2019). “Climate Change and Potential Health Effect in Benin, West Africa” International Journal of Scientific and Research Publications (IJSRP) 9(9).

Brown, J. A., M. M. Salehi, M. Moradi, B. Panahbehagh and D. R. Smith (2013). “Adaptive survey designs for sampling rare and clustered populations” Mathematics and Computers in Simulation 93: 108–116.

Case, B. K. M., J. G. Young, D. Penados, C. Monroy, L. Hebert-Dufresne and L. Stevens (2022). “Spatial epidemiology and adaptive targeted sampling to manage the Chagas disease vector Triatoma dimidiata” Plos Neglected Tropical Diseases 16(6): 18.

Chipeta, M. G., D. J. Terlouw, K. S. Phiri and P. J. Diggle (2016). “Adaptive geostatistical design and analysis for prevalence surveys” Spatial Statistics 15: 70–84.

Damien, G. B., A. Djènontin, C. Rogier, V. Corbel, S. B. Bangana, F. Chandre, M. Akogbéto, D. Kindé-Gazard, A. Massougbodji and M.-C. Henry (2010). “Malaria infection and disease in an area with pyrethroid-resistant vectors in southern Benin” Malaria Journal 9(1): 380.

Djènontin, A., S. Bio-Bangana, N. Moiroux, M.-C. Henry, O. Bousari, J. Chabi, R. Ossè, S. Koudénoukpo, V. Corbel, M. Akogbéto and F. Chandre (2010). “Culicidae diversity, malaria transmission and insecticide resistance alleles in malaria vectors in Ouidah-Kpomasse-Tori district from Benin (West Africa): A pre-intervention study” Parasites & Vectors 3(1): 83.

Fick, S. E. and R. J. Hijmans (2017). “WorldClim 2: new 1-km spatial resolution climate surfaces for global land areas” International Journal of Climatology 37(12): 4302–4315.

Fischer, G., F. Nachtergaele, S. Prieler, H. Van Velthuizen, L. Verelst and D. Wiberg (2008). “Global agro-ecological zones assessment for agriculture (GAEZ 2008).” IIASA, Laxenburg, Austria and FAO, Rome, Italy 10.

Fuhg, J. N., A. Fau and U. Nackenhorst (2020). “State-of-the-Art and Comparative Review of Adaptive Sampling Methods for Kriging” Archives of Computational Methods in Engineering 28(4): 2689–2747.

Gelfand, A. E., S. K. Sahu and D. M. Holland (2012). “On the Effect of Preferential Sampling in Spatial Prediction” Environmetrics 23(7): 565–578.

Gelman, A., B. Goodrich, J. Gabry and A. Vehtari (2019). “R-squared for Bayesian Regression Models” The American Statistician 73(3): 307–309.

Huestis, D. L., A. Dao, M. Diallo, Z. L. Sanogo, D. Samake, A. S. Yaro, Y. Ousman, Y. M. Linton, A. Krishna, L. Veru, B. J. Krajacich, R. Faiman, J. Florio, J. W. Chapman, D. R. Reynolds, D. Weetman, R. Mitchell, M. J. Donnelly, E. Talamas, L. Chamorro, E. Strobach and T. Lehmann (2019). “Windborne long-distance migration of malaria mosquitoes in the Sahel” Nature 574(7778): 404–408.

Jarvis, A., H. I. Reuter, A. Nelson and E. Guevara (2008). “Hole-filled SRTM for the globe Version 4.” available from the CGIAR-CSI SRTM 90m Database (http://srtm.csi.cgiar.org) 15(25-54): 5.

Jun, C., Y. Ban and S. Li (2014). “Open access to Earth land-cover map” Nature 514(7523): 434–434.

Kabaghe, A. N., M. G. Chipeta, R. S. McCann, K. S. Phiri, M. van Vugt, W. Takken, P. Diggle and A. D. Terlouw (2017). “Adaptive geostatistical sampling enables efficient identification of malaria hotspots in repeated cross-sectional surveys in rural Malawi” Plos One 12(2).

Kabaghe, A. N., M. G. Chipeta, D. J. Terlouw, R. S. McCann, M. van Vugt, M. P. Grobusch, W. Takken and K. S. Phiri (2017). “Short-Term Changes in Anemia and Malaria Parasite Prevalence in Children under 5 Years during One Year of Repeated Cross-Sectional Surveys in Rural Malawi” American Journal of Tropical Medicine and Hygiene 97(5): 1568–1575.

Koenraadt, C. J. M. A. S., Jeroen%A Takken, Willem (2021). Innovative strategies for vector control.

Kruschke, J. K. (2013). “Bayesian estimation supersedes the t test” J Exp Psychol Gen 142(2): 573–603.

Lazaro, E., M. Sese, A. Lopez-Quilez, D. Conesa, V. Dalmau, A. Ferrer and A. Vicent (2021). “Tracking the outbreak: an optimized sequential adaptive strategy for Xylella fastidiosa delimiting surveys” Biological Invasions 23(10): 3243–3261.

Liu, J. and J. Vanhatalo (2020). “Bayesian model based spatiotemporal survey designs and partially observed log Gaussian Cox process.” Spatial Statistics 35.

Liu, M. T., Y. Liu, L. Po, S. Xia, R. Huy, X. N. Zhou and J. M. Liu (2023). “Assessing the spatiotemporal malaria transmission intensity with heterogeneous risk factors: A modeling study in Cambodia” Infectious Disease Modelling 8(1): 253–269.

Obsomer, V., N. Titeux, C. Vancustem, G. Duveiller, J.-F. Pekel, S. Connor, P. Ceccato and M. Coosemans (2013). From Anopheles to Spatial Surveillance: A Roadmap Through a Multidisciplinary Challenge. Anopheles mosquitoes - New insights into malaria vectors.

Rebollo, M. P., H. Zoure, K. Ogoussan, Y. Sodahlon, E. A. Ottesen and P. T. Cantey (2018). “Onchocerciasis: shifting the target from control to elimination requires a new first-step-elimination mapping.” Int Health 10(suppl_1): i14–i19.

Rocha, A. D., T. A. Groen, A. K. Skidmore and L. Willemen (2021). “Role of Sampling Design When Predicting Spatially Dependent Ecological Data With Remote Sensing” IEEE Transactions on Geoscience and Remote Sensing 59(1): 663–674.

Sedda, L., E. R. Lucas, L. S. Djogbenou, A. V. C. Edi, A. Egyir-Yawson, B. I. Kabula, J. Midega, E. Ochomo, D. Weetman and M. J. Donnelly (2019). “Improved spatial ecological sampling using open data and standardization: an example from malaria mosquito surveillance” Journal of the Royal Society Interface 16(153).

Seddon, A. W. R., M. Macias-Fauria, P. R. Long, D. Benz and K. J. Willis (2016). “Sensitivity of global terrestrial ecosystems to climate variability” Nature 531(7593): 229–232.

Shrestha, H., K. McCulloch, S. M. Hedtke and W. N. Grant (2022). “Geospatial modeling of pre-intervention nodule prevalence of Onchocerca volvulus in Ethiopia as an aid to onchocerciasis elimination” Plos Neglected Tropical Diseases 16(7): 25.

Stolk, W. A., D. J. Blok, J. I. D. Hamley, P. T. Cantey, S. J. de Vlas, M. Walker and M. G. Basanez (2021). “Scaling-Down Mass Ivermectin Treatment for Onchocerciasis Elimination: Modeling the Impact of the Geographical Unit for Decision Making.” Clinical Infectious Diseases 72: S165–S171.

Thawer, S. G., M. Golumbeanu, K. Munisi, S. Aaron, F. Chacky, S. Lazaro, A. Mohamed, N. Kisoka, C. Lengeler, F. Molteni, A. Ross, R. W. Snow and E. Pothin (2022). “The use of routine health facility data for micro-stratification of malaria risk in mainland Tanzania” Malaria Journal 21(1): 14.

Wang, Y. K., C. L. Moe, S. Dutta, A. Wadhwa, S. Kanungo, W. Mairinger, Y. C. Zhao, Y. Jiang and P. F. M. Teunis (2020). “Designing a typhoid environmental surveillance study: A simulation model for optimum sampling site allocation.” Epidemics 31.

World Health Organization and UNICEF (2017). “Global vector control response 2017-2030”

